# Inhibition of EZH2 ameliorates lupus-like disease in *MRL/lpr* mice

**DOI:** 10.1101/500918

**Authors:** Dallas M. Rohraff, Ye He, Evan A. Farkash, Mark Schonfeld, Pei-Suen Tsou, Amr H. Sawalha

## Abstract

**Objectives:** We previously revealed a role for EZH2 in inducing pro-inflammatory epigenetic changes in lupus CD4+ T cells. In this study, we sought to determine if inhibiting EZH2 ameliorates lupus-like disease in MRL/*lpr* mice.

**Methods:** EZH2 expression levels in multiple cell types in lupus patients were evaluated using flow cytometry and mRNA expression data. Inhibition of EZH2 in MRL/*lpr* mice was achieved by DZNep intraperitoneal administration using a preventative and a therapeutic treatment model. Effects of DZNep on animal survival, anti-dsDNA antibody production, proteinuria, renal histopathology, cytokine production, and T and B cell numbers and percentages were assessed.

**Results:** EZH2 expression levels were increased in whole blood, neutrophils, monocytes, B cells, and CD4+ T cells in lupus patients. In MRL/*lpr* mice, inhibiting EZH2 with DZNep treatment before or after disease onset improved survival and significantly reduced anti-dsDNA antibody production. DZNep-treated mice displayed a significant reduction in renal involvement, splenomegaly, and lymphadenopathy.

Lymphoproliferation and numbers of double-negative T cells were significantly reduced in DZNep treated mice. Concentrations of circulating cytokines and chemokines, including TNF, IFN-γ, CCL2, RANTES/CCL5, IL-10, KC/CXCL1, IL-12, IL-12p40 and MIP-1β/CCL4 were decreased in DZNep treated mice.

**Conclusions:** ZH2 is upregulated in multiple cell types in lupus patients. Therapeutic inhibition of EZH2 abrogates lupus-like disease in MRL/*lpr* mice, suggesting that EZH2 inhibitors may be repurposed as a novel therapeutic option in lupus patients.

## Introduction

Systemic Lupus Erythematosus (SLE or lupus) is a chronic relapsing autoimmune disease that involves multiple organ systems. Lupus is characterized by the production of autoantibodies directed against nuclear antigens, and a dysregulated immune response. The etiology of lupus remains unknown, however, both genetic and epigenetic mechanisms have been shown to contribute to disease pathogenesis (1, 2).

DNA methylation plays a critical role in the pathogenesis of lupus (3). Abnormal DNA methylation patterns have been described in multiple immune cell types isolated from lupus patients, and a role for genetic-epigenetic interaction in the pathogenesis of lupus has been suggested (4). Further, abnormal DNA methylation patterns in lupus have been shown to contribute to clinical heterogeneity, disease variability between ethnicities, and lupus flare and remission (4).

We have recently demonstrated that increased disease activity in lupus patients is characterized by an early epigenetic shift in naïve CD4+ T cells the precedes CD4+ T cell differentiation and effector T cell transcriptional activity (5). We provided evidence that this epigenetic shift is likely induced by EZH2 overexpression as a result of downregulation of miR-101 and miR-26a, and demonstrated that EZH2 overexpression mediates increased T cell adhesion in lupus due to EZH2-induced demethylation and transcriptional de-repression of the adhesion molecule JAM-A (5, 6).

EZH2 overexpression has been linked to increased invasiveness in a number of malignancies, and EZH2 inhibitors are currently being evaluated in clinical trials for cancer therapy (7). Our data indicate that inhibiting EZH2 might be of therapeutic potential in lupus, suggesting the possibility of pharmacologic repurposing of EHZ2 inhibitors as a therapeutic option for lupus. In this study, we first examined EZH2 expression patterns in other immune cell types isolated from the peripheral blood of lupus patients, and then tested the effects of using an EZH2 inhibitor in the lupus-prone MRL/*lpr* mouse model, utilizing both preventative and therapeutic approaches. Our data provide robust pre-clinical evidence supporting the potential use of EZH2 inhibitors in lupus, paving the way for repurposing EZH2 inhibitors in lupus clinical trials.

## Materials and Methods

### EZH2 expression in B cells, monocytes, and neutrophils in lupus patients

We recruited a total of six lupus patients (mean ± SEM age: 46.2 ± 5.4; age range: 32 to 61 years) and six healthy controls (mean ± SEM age: 44.5 ± 5.8; age range: 30 to 63 years). All lupus patients fulfilled the American College of Rheumatology classification criteria for SLE (8). The mean SLEDAI score for lupus patients was 2 with a median of 1 (range: 0 to 6). Lupus patients on cyclophosphamide or methotrexate were excluded from participating in the study. All participants signed informed consent approved by the institutional review board of the University of Michigan. Peripheral whole blood was collected from each study subject, and erythrocytes in blood samples were lysed with RBC Lysis Buffer (eBioscience). Lysed whole blood cells were incubated with Human Seroblock (BIO-RAD) in cell staining buffer (BioLegend) to block unspecific Fc receptor binding of Ig, and extracellularly stained with the following fluorochrome-conjugated antibodies: APC/Cy7 anti-CD14 (clone: 63D3, BioLegend), PE Mouse Anti-Human CD16b (clone: CLB-gran11.5 (RUO), BD Biosciences), APC anti-CD19 (clone: HIB19, BioLegend). Afterward, cells were fixed with Fixation Buffer (BioLegend), permeabilized with Intracellular Staining Permeabilization Wash Buffer (BioLegend), and subsequently stained with FITC anti-EZH2 (clone: REA907, Miltenyi Biotec) or with corresponding isotype control, REA Control (I) antibody (clone: REA293, Miltenyi Biotec). Stained cells were then analyzed by flow cytometry using an iCyt Synergy SY3200 Cell Sorter (Sony Biotechnology Inc.) and by FlowJo software v10.0.7 (Treestar). An example of the gating strategy to determine B cells, monocytes, and neutrophils is shown in **Supplementary Figure 1**, based on FSC versus SSC and positive expression of their respective markers. The expression of EZH2 is measured by median fluorescence intensity (MFI) of EZH2 antibody minus MFI of its isotype control.

### GEO data analysis

Analysis of EZH2 mRNA expression levels between lupus patients and healthy controls (in whole blood cells, CD4+ T cells, and CD19+ B cells) was performed using datasets downloaded from Gene Expression Omnibus (GEO) (accession numbers GSE72509 and GDS4185). These datasets have been described in detail previously (9, 10).

### Mice and Treatments

Eight-week-old female MRL/*lpr* mice (#000485; The Jackson Laboratory) were acclimatized for two weeks prior to study commencement and maintained under pathogen-free conditions. DZNep (Cayman Chemical) was administered in two model types: a preventative model with DZNep treatments beginning at 10 weeks of age (DZNep/DZNep, Day 0 of the study) and a therapeutic treatment model with DZNep treatments beginning at 14 weeks of age (Vehicle/DZNep, Day 28 of the study). A vehicle group was also included as control. Mice were treated with either vehicle or DZNep (2 mg/kg) through intraperitoneal injection. DZNep was solubilized in DMSO and diluted in PBS prior to injection (final DMSO concentration was 10% in PBS). For the Vehicle control group, mice received daily injection of Vehicle control for 35 days and switched to twice a week (Monday and Thursday) dosing until day 98. A similar regimen was used for the DZNep/DZNep prevention model group; mice received DZNep once daily from Day 0 to Day 35 and then switched to twice weekly dosing for the remainder of the study. For the Vehicle/DZNep therapeutic treatment group, animals received vehicle once daily from Day 0 to Day 27. On Day 28, mice were switched to a once daily dosage of DZNep until Day 63; from Day 64 onwards, the dosing frequency was changed to twice weekly. Animals were monitored and weighed daily throughout the study. This study was performed at Biomodels, an AAALAC accredited facility, and approved by the Biomodels IACUC.

### Assessment of lupus nephritis and kidney damage

Kidneys were harvested on Day 98 of the study, fixed in formalin, and sections were cut and stained with either hematoxylin and eosin or Periodic acid-Schiff stain. Clinical scores assessing glomerulonephritis, crescent formation, and necrosis in kidneys were obtained in a blinded manner, without knowledge of the experimental group. Approximately 125 glomeruli in each periodic acid Schiff stained slide were counted by a pathologist blinded to treatment. Glomeruli were categorized as having crescents, fibrinoid necrosis, acute glomerulitis, mesangial hypercellularity, segmental glomerulosclerosis, global glomerulosclerosis, or normal architecture. The presence of pseudothrombi, interstitial nephritis, and arteritis was also noted.

Urine was collected every two weeks beginning two weeks prior to Day 0 of the study. Urine albumin and creatinine concentrations were measured using enzyme-linked immunosorbent assay (ELISA) kits from Alpha Diagnostic and R& D system, respectively, and used to calculate the urine albumin-to-creatinine ratio (UACR).

### Quantification of serum and plasma antibodies and cytokines

Every two weeks, serum was collected by retro orbital bleed using serum separator tubes. On Day 98 of the study, plasma was collected from all the animals while serum was collected in a portion of the DZNep/DZNep-treated mice and all the Vehicle/DZNep-treated group. Anti-double-stranded DNA (anti ds-DNA) antibody titers in serum or plasma (day 98 only) were determined using ELISA (Alpha Diagnostic). Cytokines in plasma collected on day 98 were quantified using the Bio-Plex ProTM Mouse Cytokine 23-plex Assay (Bio-Rad).

### Spleen and lymph node weights

Spleen and lymph nodes (submaxillary, thoracic, axillary, renal, and mesenteric) were excised on day 98 of the study, trimmed of extra fat and connective tissue, weighed, and photographed. The average weight of all lymph nodes in each mouse was used to represent the average change in lymph node weight between mouse groups.

### Flow Cytometric analysis of Spleen Cells

Each spleen was placed in FACS buffer (0.5% BSA, 2mM EDTA in PBS) and processed into single cell suspension using a gentleMACSTM Dissociator. Total spleen cell counts were obtained by counting of a fraction of the diluted cells by flow cytometry. Red blood cells (RBC) were lysed using BD Pharm Lyse lysis buffer (BD Biosciences) prior to the addition of mouse FcR Blocking Reagent (Miltenyi Biotec). Cells were stained with the following fluorochrome-conjugated antibodies: CD3ε-VioBlue (clone: 17A2, Miltenyi Biotec), APC-Cy™7 Rat Anti-Mouse CD4 (clone: GK1.5 (RUO), BD Biosciences), PE Rat Anti-Mouse CD8a (clone: 53-6.7 (RUO), BD Biosciences), PerCP/Cy5.5 anti-mouse TCR β chain (clone: H57-597, BioLegend), FITC anti-mouse CD19 (clone: 6D5, BioLegend), CD45R (B220)-APC, (clone: RA3-6B2, Miltenyi Biotec), and PE/Cy7 anti-mouse CD11c (clone: N418, BioLegend). Stained cells were then analyzed by flow cytometry using a MACSQuant^®^ Analyzer 10 (Miltenyi Biotec) and with WinList Version 9 software (Verity Software House). The following cell types were gated based on FSC versus SSC and positive expression of their respective markers: B cells (TCRβ-CD19+ cells), total T cells (TCRβ+ cells), CD4+ T cells (TCRβ+CD4+ cells), CD8+ T cells (TCRβ+CD8+ cells), and double-negative (DN) T cells (TCRβ+CD4-CD8-cells). An example of the gating strategy is shown in **Supplementary Figure 2**.

### Cytokine Analysis

Cytokine levels in plasma from day 98 of the mouse study were analyzed using a Bio-Plex ProTM Mouse Cytokine 23-plex Assay (Bio-Rad). The cytokines assayed include TNF, IFN-γ, IL1α, IL-1β, IL-2, IL-3, IL-4, IL-5, IL-6, IL-9, IL-10, IL-12, IL-12p40, IL-13, IL-17a, CCL2, RANTES/CCL5, KC/CXCL1, Exotaxin/CCL11, MIP-1α, MIP-1β/CCL4, G-CSF, and GM-CSF.

### Statistical Analysis

Mann-Whitney U tests were used to compare the publicly available data of human EZH2 expression levels in whole blood, B cells, and CD4+ T cells in lupus patients and healthy controls. Comparison of EZH2 expression between lupus patients and controls in B cells, monocytes, and neutrophils by flow cytometry was compared using an Unpaired t test or a Mann-Whitney U test when normality could not be assumed. Survival Analysis was evaluated using the Mantel-Cox log-rank test.

All other mouse data were evaluated using a one-way ANOVA, or a Kruskal-Wallis test when normality could not be assumed, with Dunn’s multiple comparison test to compare the DZNep/DZNep and Vehicle/DZNep groups to the Vehicle control. All statistical analyses were performed using GraphPad Prism version 7 (GraphPad Software, Inc.). *P*-values of less than 0.05 were considered statistically significant. Data are presented as mean ± SEM.

## Results

### EZH2 expression in lupus

We previously showed that EZH2 is upregulated in lupus CD4+ T cells, and that T cell overexpression of EZH2 plays an important role in lupus (5, 6). To examine if other immune cell types also overexpress EZH2 in lupus, we determined EZH2 expression using flow cytometry in B cells, monocytes, and neutrophils isolated from peripheral blood from lupus patients compared to normal healthy controls. EZH2 expression, as determined by median fluorescence intensity (MFI), was significantly elevated in B cells, monocytes, and neutrophils of lupus patients compared to matched controls (p<0.05; **Figures 1A, 1B**, and **1C**). Using previously published gene expression profiles available in Gene Expression Omnibus (GEO), we analyzed EZH2 mRNA levels in whole blood (GEO accession GSE72509) and freshly isolated lymphocyte subsets (GEO Accession GDS4185) of lupus patients and healthy controls. EZH2 mRNA levels in whole blood of lupus patients were significantly higher than healthy controls (p<0.01; **Figure 1D**). Similarly, EZH2 expression was significantly elevated in the freshly isolated B cells and CD4+ T cells of lupus patients when compared to controls (p<0.05; **Figures 1E** and**1F**).

**Figure 1.**
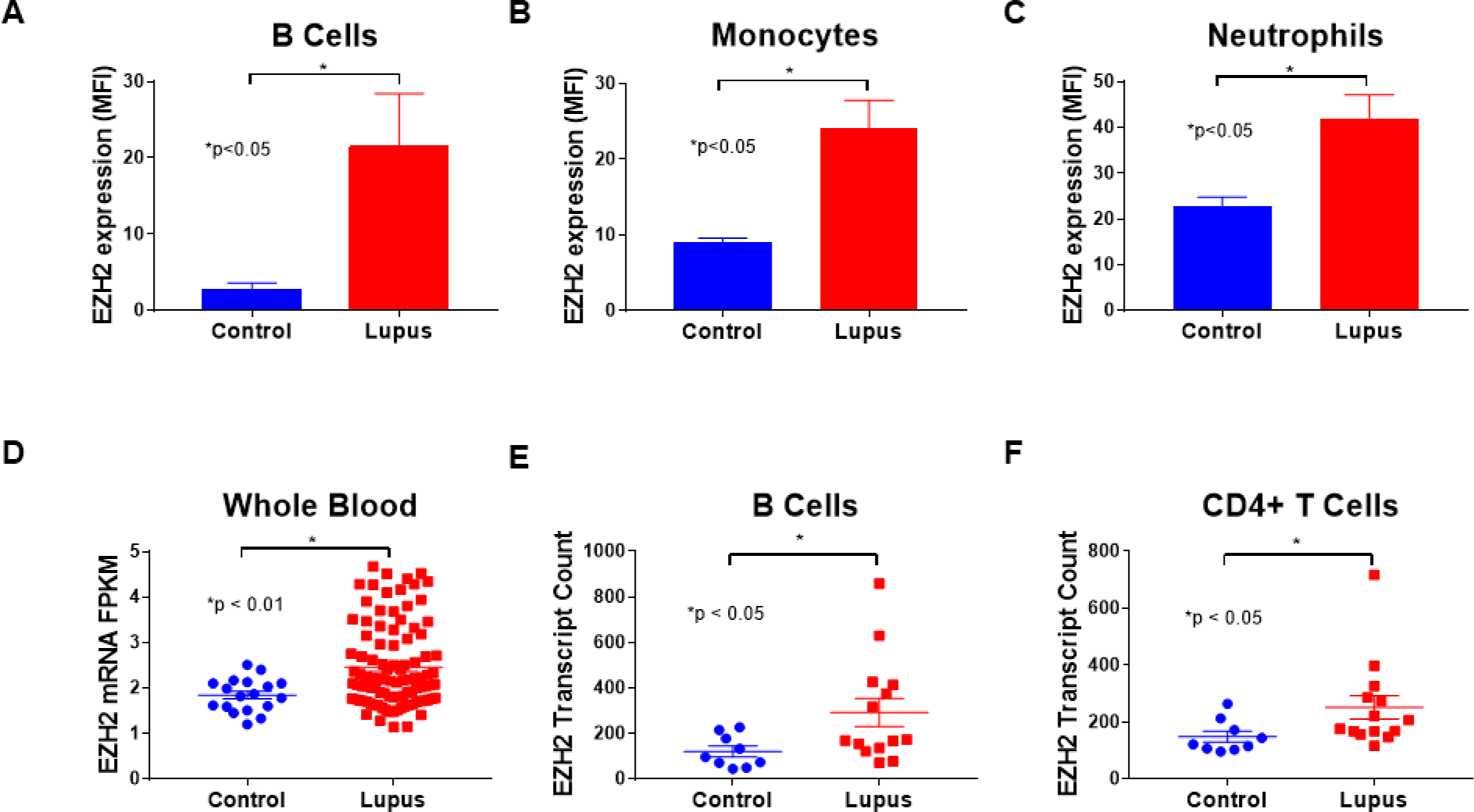
Analysis of EZH2 expression in human peripheral blood cells. EZH2 expression, analyzed by flow cytometry, was elevated in B cells **(A)**, monocytes **(B)**, and neutrophils **(C)** of lupus patients (n=6) versus controls (n=6). **(D)** EZH2 mRNA levels were elevated in the whole blood of lupus patients (n=99) compared to controls (n=18). EZH2 mRNA levels were also elevated in the B cells **(E)** and CD4+ T cells **(F)** of lupus patients (n=14) compared to controls (n=9). Results are expressed as mean +/-SEM and p<0.05 was considered significant.

### Effect of EZH2 inhibition on mortality and autoantibody production in MRL/*lpr* mice

To examine if inhibition of EZH2 is beneficial in lupus-like disease, an EZH2 inhibitor, DZNep, was administered to MRL/*lpr* mice (**Figure 2A** and **2B**). As shown in **Figure 2C**, mice that received only vehicle had 33.3% mortality by the end of the study at Day 98 (mice were at 24 weeks of age). Mice in the preventative group (DZNep/DZNep) had 100% survival, while there was 6.67% mortality in the therapeutic group (Vehicle/DZNep) by the end of the study. There was a significant difference in the mortality rates between the DZNep/DZNep-treated and the Vehicle control group (p<0.05).

There were significantly lower anti-dsDNA antibody levels in DZNep/DZNep group by Day 14, two weeks after DZNep treatment began in this group, and anti-dsDNA antibody levels remained significantly lower throughout the duration of the study (p<0.001, **Figure 2D**). By Day 56, the anti-dsDNA titers were significantly lower in the Vehicle/DZNep group compared to Vehicle control (p<0.05) and remained significantly lower for the next 28 days, though there was not a significant difference detected at the final timepoint (**Figure 2D**). Although plasma, instead of serum, was used for quantification of anti-dsDNA at Day 98, the levels were similar to what was measured in serum (data not shown). Our results showed that DZNep improved survival in the MRL/*lpr* mice and significantly reduced the production of anti-dsDNA antibodies two to four weeks following daily dosing of DZNep.

**Figure 2.**
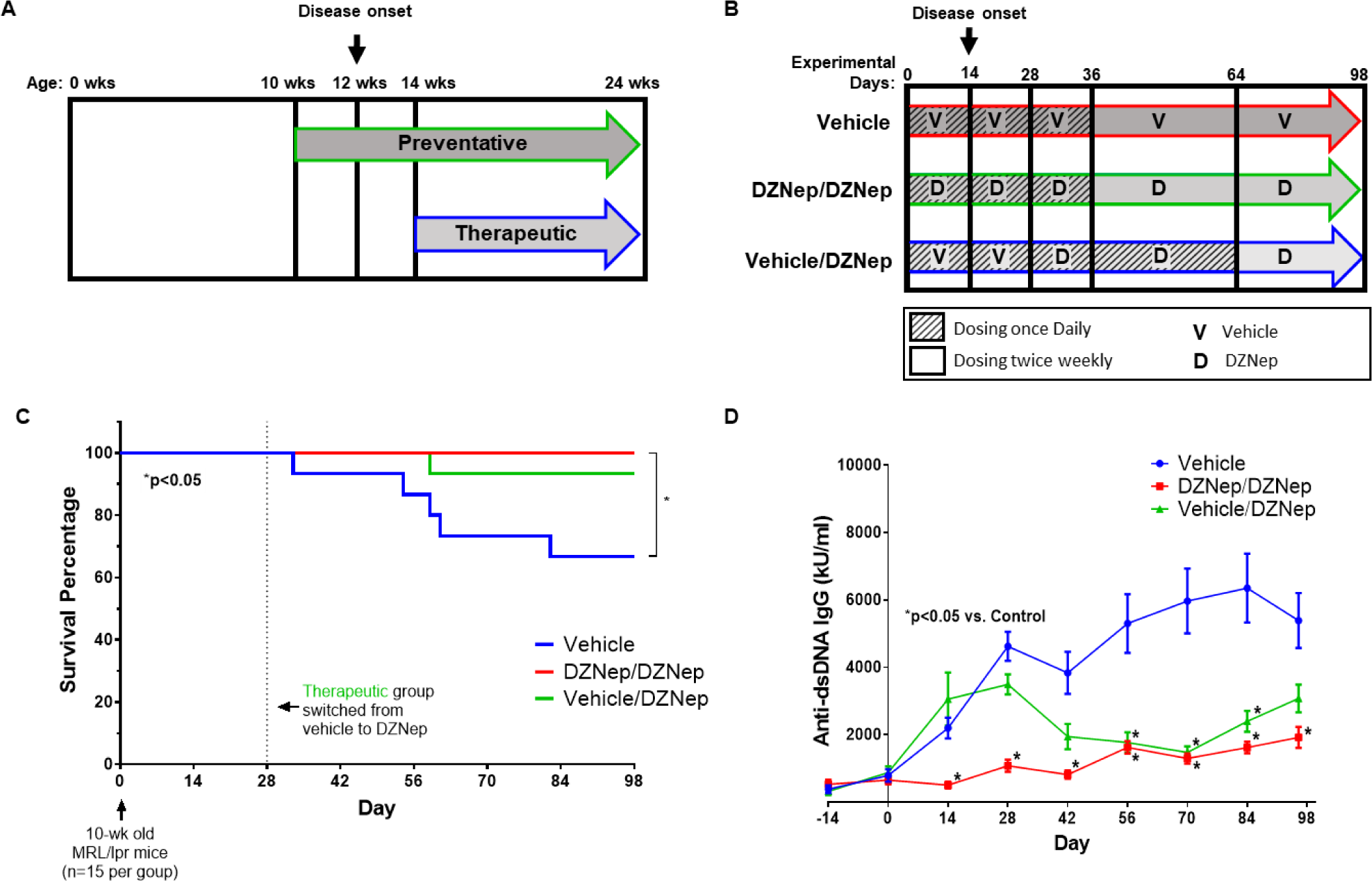
DZNep treatment in MRL/*lpr* mice improves survival and reduces anti-dsDNA antibody production. **(A)** Schematic representation of preventative and therapeutic treatment models. DZNep treatment in the preventative (DZNep/DZNep) group began when mice were 10 weeks old; two weeks prior to disease onset. The therapeutic group (Vehicle/DZNep) received DZNep treatment two weeks after disease onset when mice were 14 weeks old. **(B)** Diagram of treatment regimens. The DZNep/DZNep group received DZNep once daily for 35 days, and then dosing of DZNep was switched on Day 36 to twice weekly until the end of the study. The Vehicle control group followed the same regimen as the DZNep/DZNep group with Vehicle administered instead of DZNep. The Vehicle/DZNep group received Vehicle once daily for 27 days, then switched to daily dosing of DZNep from Day 28 until Day 63. On Day 64, dosing of DZNep was reduced to twice weekly until the end of the study. **(C)** Survival curve of the Vehicle, DZNep/DZNep, and Vehicle/DZNep groups (n=15 MRL/*lpr* mice per group at start). **(D)** Anti-dsDNA antibody levels were monitored beginning 14 days before Day 0 through Day 98 of the study. Results are expressed as mean +/-SEM and p<0.05 was considered significant.

### Effect of DZNep treatment on renal involvement

To assess the effect of DZNep on renal damage, glomerulonephritis, crescent formation, and necrosis were assessed for each mouse on Day 98 of the study. Development of glomerulonephritis and crescents were significantly reduced in both the DZNep/DZNep and Vehicle/DZNep groups compared to Vehicle control (p<0.05, **Figures 3A** and **3B**). There also was a significant reduction in glomerular necrosis in the DZNep/DZNep group compared to Vehicle control (p<0.01), but the difference between the Vehicle and Vehicle/DZNep group was not statistically significant (**Figure 3C**). Overall, the number of glomeruli with no pathologic abnormality was significantly higher in the DZNep/DZNep group compared to Vehicle control (65.8%±4.7 versus 31.6%±7.2 [mean±SEM], p<0.01; **Supplemental Figure 3**). To monitor the progression of kidney involvement, proteinuria was analyzed using urine albumin-to-creatinine ratios (UACR) calculated every two weeks throughout the study. UACR were significantly lower in the DZNep/DZNep and Vehicle/DZNep groups compared to Vehicle controls by Day 14 and 42, respectively (p<0.01, **Figure 3E**). For both treatment groups, UACR remained significantly lower throughout the remainder of the study. Overall, DZNep treatment, both preventative and therapeutic, significantly reduced lupus nephritis and renal damage in MRL/*lpr* lupus-prone mice.

**Figure 3.**
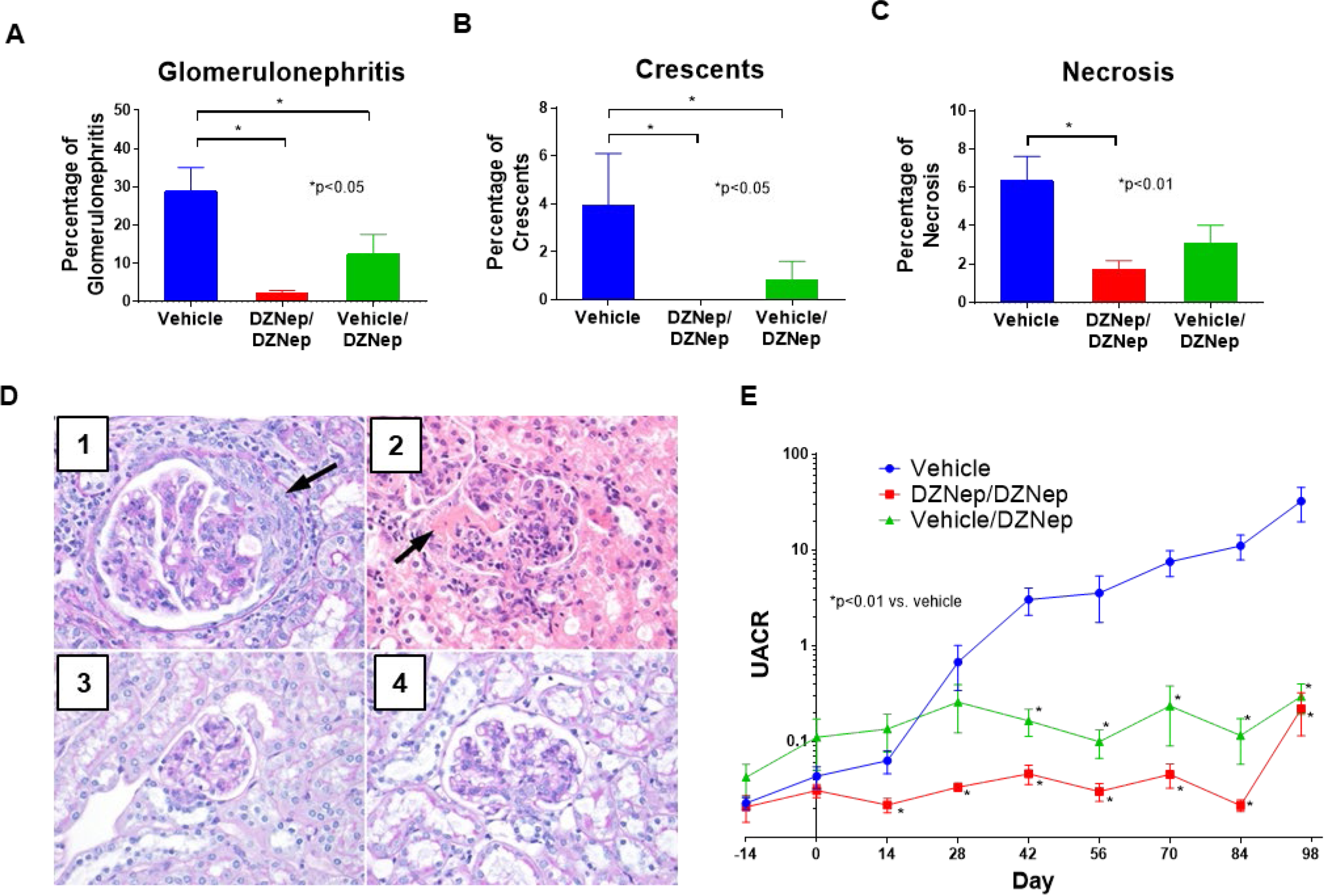
DZNep treatment reduced renal involvement in MRL/*lpr* mice. The percentage of total glomeruli with glomerulonephritis **(A)**, crescent formation **(B)**, and necrosis **(C)** in the kidneys of the Vehicle, DZNep/DZNep, and Vehicle/DZNep groups. **(D)** Photographic representation of renal damage in kidney histopathology. **(D1)** Glomerulus with glomerulomegaly and cellular crescent (arrow) from a Vehicle control mouse (Periodic acid Schiff stain, 400x). **(D2)** Acute global glomerulitis and segmental fibrinoid necrosis (arrow) from a Vehicle control mouse (Hematoxylin & Eosin, 400x). **(D3)** Normal appearing glomerulus from mouse in DZNep/DZNep group (Periodic acid Schiff stain, 400x). **(D4)** Glomerulus with mesangial hypercellularity from mouse in Vehicle/DZNep group (Periodic acid Schiff stain, 400x). **(E)** Urine albumin-to-creatinine ratios (UACR) in the Vehicle, DZNep/DZNep, and Vehicle/DZNep groups. Results are expressed as mean +/-SEM and p<0.05 was considered significant.

### Reduced splenomegaly and lymphadenopathy with DZNep

MRL/*lpr* mice develop progressive enlargement of the spleen and lymph nodes due to lymphoproliferation that is characteristic of the strain. The spleen from Vehicle-treated mice weighed significantly more than that of both the DZNep/DZNep-treated and the Vehicle/DZNep-treated groups (p<0.001; **Figure 4A**). The mean weight of lymph nodes from all sites (submaxillary, thoracic, axillary, renal, and mesenteric) in both the DZNep/DZNep and the Vehicle/DZNep groups was significantly lower than the Vehicle control group (p<0.0001; **Figure 4B, Supplemental Figure 4**). DZNep appeared to be efficacious in reducing the progression of splenomegaly and lymphadenopathy in MRL/*lpr* mice.

**Figure 4.**
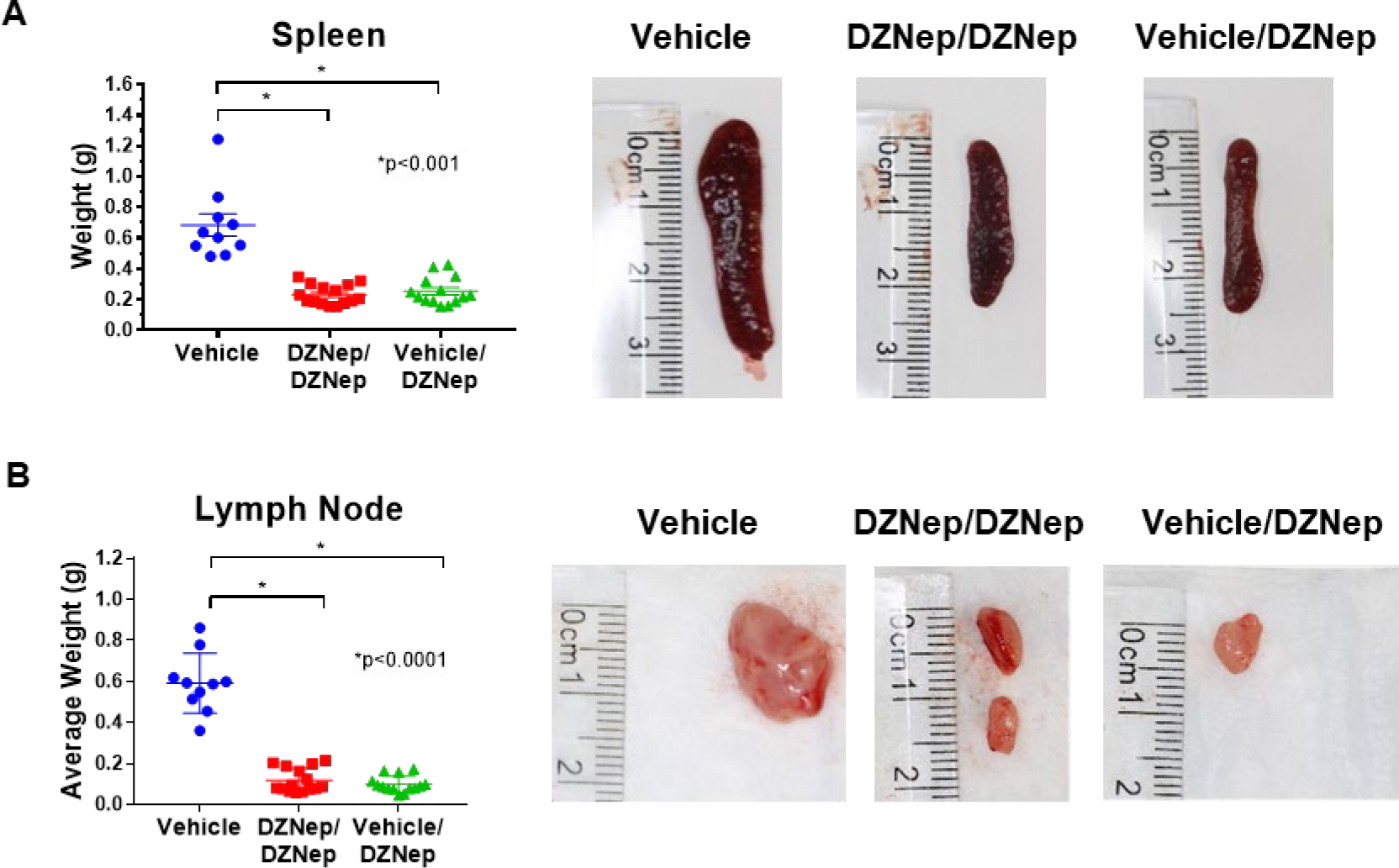
Splenomegaly and lymphadenopathy are reduced with DZNep treatment in MRL/*lpr* mice. **(A)** The weight of each mouse spleen and representative photographs of a mouse spleen from each treatment group. **(B)** The average weight of all lymph node groups per mouse and representative lymph node photographs (renal lymph node of median weight from each treatment group). Results are expressed as mean +/-SEM and p<0.05 was considered significant.

### Reduced lymphoproliferation with DZNep treatment

We further explored the effect of EZH2 inhibition on lymphoproliferation in MRL/*lpr* mice by analyzing differences in T and B cell populations between mouse groups using flow cytometry analysis. Both the DZNep/DZNep and the Vehicle/DZNep-treated groups had reduced total numbers of splenocytes, total T cells (TCRβ+ cells), CD8+ T cells (CD8+TCRβ+ cells), and DN T cells (CD4-CD8-TCRβ+ cells) when compared to Vehicle control mice (p<0.01; **Figures 5A**, **5B**, and **5C**). The total number of CD4+ T cells (CD4+TCRβ+ cells) was significantly decreased in the Vehicle/DZNep group compared to control (p<0.01; **Figure 5C**). In addition, DZNep treatment in both the DZNep/DZNep and the Vehicle/DZNep-treated groups significantly reduced the percentage of DN T cells while significantly increasing the percentage of CD4+ and CD8+ T cells at the end of the study when compared to the Vehicle control group (p≤0.0001; **Figure 5D**). The observed shift in T cell populations and reduction in total number of T cells suggests DZNep reduces T cell hyperproliferation and reduces the generation of pathogenic DN T cells in MRL/*lpr* mice.

**Figure 5.**
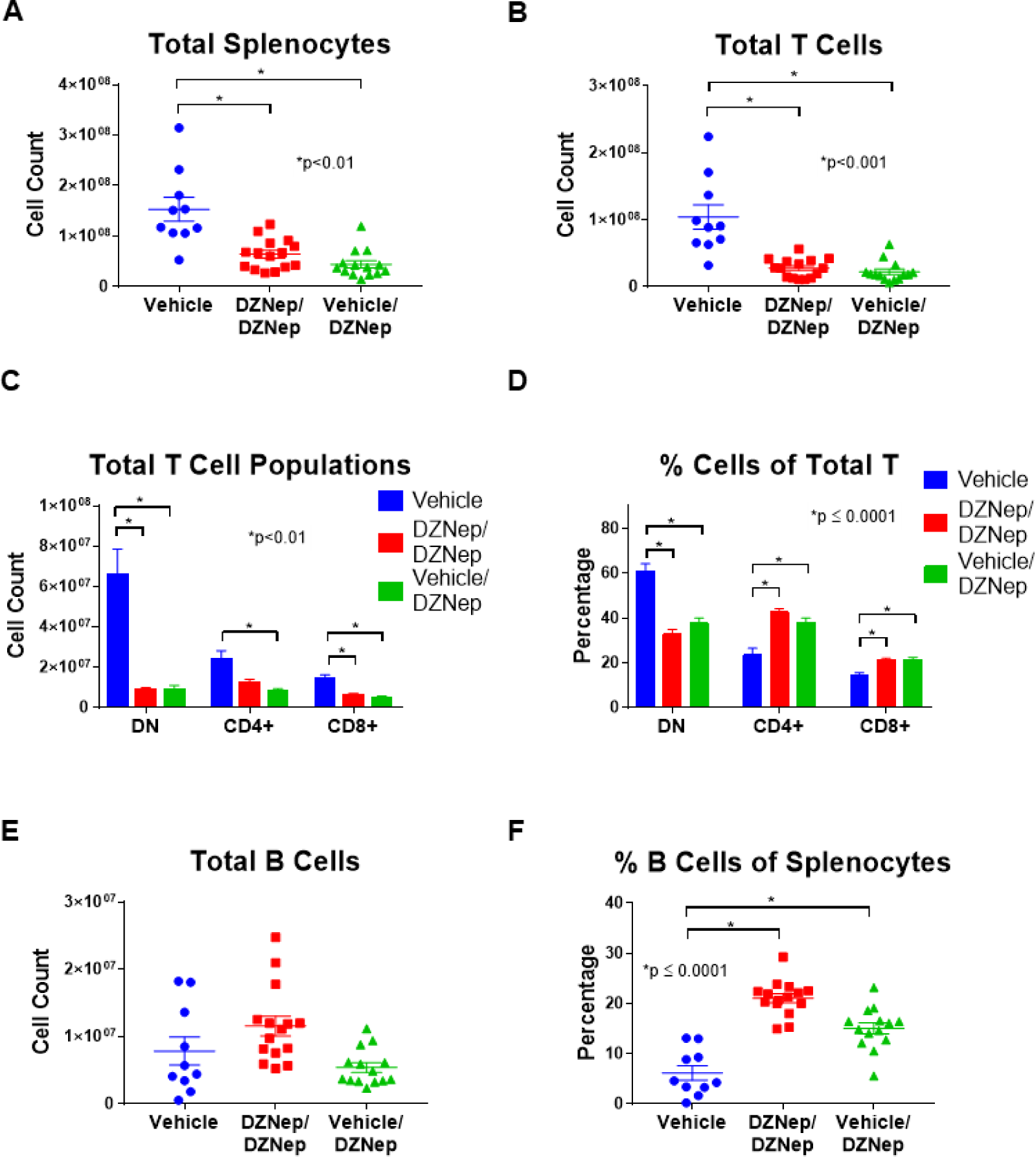
Reduced lymphoproliferation is observed in DZNep treated MRL/*lpr* mice. DZNep treatments significantly reduced the total number of splenocytes **(A)** and total T cells (TCRβ+ cells) **(B)** in DZNep/DZNep and Vehicle/DZNep compared to Vehicle controls. Total numbers **(C)** and percentages **(D)** of DN T cells (TCRβ+CD4-CD8-cells), CD4+ T cells (TCRβ+CD4+ cells), and CD8+ T cells (TCRβ+CD8+ cells) in the Vehicle, DZNep/DZNep, and Vehicle/DZNep groups. Total numbers of B cells **(E)** and percentages **(F)** in DZNep treatment groups compared to Vehicle control. Results are expressed as mean +/-SEM and p<0.05 was considered significant.

There was no difference in the total number of B cells (CD19+TCRβ-cells) between any groups (**Figure 5E**). We observed, however that the B cell percentage of total splenocytes (CD19+TCRβ-cells of total splenocytes) was significantly greater in both the DZNep/DZNep and Vehicle/DZNep groups when compared to controls (p≤0.0001; **Figure 5F**).

### Cytokine analysis

To further characterize the effect of EZH2 inhibition in MRL/*lpr* mice, we analyzed the differential expression of plasma cytokine levels at day 98. Cytokine levels were undetectable for IL-3, IL-1b, IL-9, IL-13, IL-6, IL-4, G-CSF, GM-CSF, and MIP-1a. No significant differences were observed between DZNep treatment groups and vehicle controls for IL-2, IL-5, IL1a, IL-17a, and Exotaxin/CCL11 (data not shown). However, there was a significant reduction in plasma levels of TNF, IFN-γ, CCL2, RANTES/CCL5, IL-10, KC/CXCL1, IL-12, IL-12p40 and MIP-1β/CCL4 in both the DZNep/DZNep-treated and Vehicle/DZNep-treated groups when compared to the Vehicle control group (**Figure 6)**.

**Figure 6.**
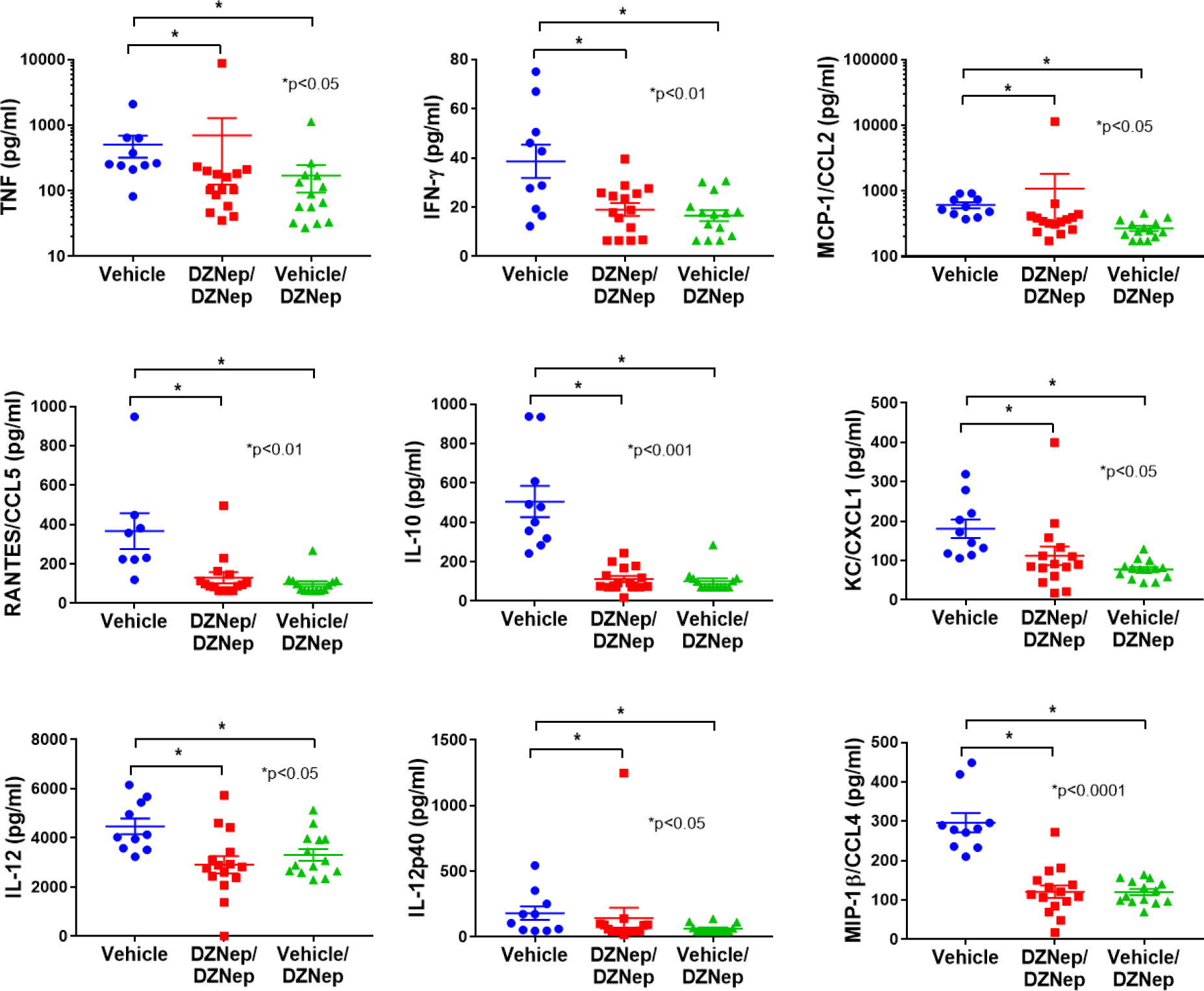
Decreased levels of plasma cytokines with DZNep treatment. There was a significant reduction in plasma levels of TNF, IFN-γ, MCP-1/CCL2, RANTES/CCL5, IL-10, KC/CXCL1, IL-12, IL-12p40 and MIP-1β/CCL4 in both the DZNep/DZNep-treated and Vehicle/DZNep-treated groups when compared to the Vehicle control group. Results are expressed as mean +/-SEM and p<0.05 was considered significant.

## Discussion

Our previous work suggested that EZH2 plays an important role in lupus (5, 6). We reported that naïve CD4+ T cells from lupus patients had higher expression levels of EZH2 than healthy controls, and this upregulation is critical for T cell adhesion, as inhibition of EZH2 by DZNep can normalize the ability of lupus CD4+ T cells to adhere to endothelial cells (6). In this study we further examined the expression of EZH2 in other cell types and showed that EZH2 was significantly upregulated in B cells, monocytes, and neutrophils in lupus patients. These data prompted us to investigate whether inhibition of EZH2 would be beneficial for lupus. Indeed, in MRL/*lpr* mice, DZNep treatment, either dosing following a preventative or a therapeutic fashion, improved survival and reduced autoantibody production. In addition, the therapeutic effects of DZNep appear rapidly, as lowering anti-dsDNA antibody levels and UACR were evident 14 days following the first dose of DZNep.

The improvement in survival observed in DZNep-treated mice is likely in part due to a reduction in renal involvement. DZNep treatments appeared to prevent the progression of renal damage, as evidenced by relatively steady UACR in DZNep-treated mice. Renal damage was also assessed at the end of the study through histopathological examination. We observed a significant reduction in glomerulonephritis and crescent formation in both DZNep treatment groups, and a significant reduction in glomerular necrosis in the DZNep/DZNep group compared to Vehicle control by the end of the study. The development of lupus nephritis is indicative of severe disease in lupus patients and is an important predictor of morbidity and mortality. Because DZNep treatment effectively reduced renal involvement in MRL/*lpr* mice, an EZH2 inhibitor could similarly prove therapeutic for lupus patients.

The MRL/*lpr* mouse model is a lymphoproliferative model characterized by T cell and B cell dysregulation (11). The infiltration of DN T cells, in addition to an overall hyperproliferation of T cell populations contribute significantly to the observed splenomegaly and lymphadenopathy in MRL/*lpr* mice (11, 12). We observed a significant reduction in spleen and lymph node weights in both groups of mice treated with DZNep. A significant reduction in the total number of splenocytes in DZNep-treated groups was also observed, suggesting a reduction in lymphoproliferation. While DZNep treatment in both groups reduced the total number of T cells, CD4+ T cells, and CD8+ T cells, the reduction in CD4-CD8-T cells (DN T cells) is of particular interest. DN T cells represent a pathogenic T cell subset in lupus likely derived from CD8+ T cells (13). DN T cells contribute to the production of autoantibodies, produce significant amounts of inflammatory cytokines including IL-17 and IFN-γ, and contribute to renal damage as they accumulate in the kidneys of lupus patients (13-16). Inhibition of EZH2 with DZNep significantly reduced the total number of pathogenic DN T cells in MRL/*lpr* mice. Indeed, the significant reduction in DN T cell numbers observed with DZNep caused significant elevation in the percentages of the single positive T cells (CD4+ and CD8+ T cells). We speculate that the reduction of pathogenic DN T cells in the DZNep-treated groups likely contributed to the observed reduction in renal damage and levels of circulating autoantibodies.

Cytokine and chemokine production are important in lupus pathogenesis and tissue damage in lupus patients. We observed reduced concentrations of TNF, IFN-γ, CCL2, RANTES/CCL5, IL-10, IL-12, IL-12p40, and MIP-1β/CCL4 in the plasma of the DZNep-treated groups compared to controls. All of these cytokines and chemokines have been reported to be elevated in lupus patients (17-21). Chemokines such as RANTES/CCL5, MCP-1/CCL2, and MIP-1β/CCL4 contribute to lupus pathogenesis by recruiting leukocytes and other effector cells to inflamed tissues, causing damage (17-19). IL-10 and TNF have been reported to be upregulated and correlated with disease activity in lupus patients experiencing renal involvement (17, 20). IFN-γ has been shown to contribute to the development of lupus through promotion of autophagy in lupus T cells, and potentially the persistence of pathogenic T cell subsets (20). Dysregulation of IL-12 has been reported in patients with lupus (22), and a recent clinical trial adding ustekinumab, which targets IL-12p40, to standard-of-care in lupus patients in a phase II trial showed promising results (23).

We have previously demonstrated that EZH2 might mediate a pathogenic effect in lupus though altering T cell DNA methylation patterns and upregulation of the adhesion molecule JAM-A in CD4+ T cells (5, 6). As the Notch signaling pathway regulates effector T cell survival and function including cytokine production (24-26), we speculate that another potential mechanism of action for EZH2 in lupus might involve the Notch signaling pathway. EZH2 has been shown to inhibit T cell Notch repressors, promoting Notch activation and thus effector T cell polyfunctionality and survival (27). Notch pathway is also critical in B cells and monocytes (28-32), which are also characterized by EZH2 overexpression in lupus in our study.

In summary, we demonstrate that EZH2 in overexpressed in multiple immune cell types in lupus patients. Inhibiting EZH2 is associated with abrogation of lupus-like disease in MRL/*lpr* mice. Our data suggest that inhibiting EZH2 might provide a novel therapeutic approach for lupus patients.

## Conflict of interest

The authors declare no conflicts of interest.

## Funding

This work was supported by the Lupus Research Alliance and the National Institute of Allergy and Infectious Diseases of the National Institutes of Health grant number R01AI097134. Dr. Farkash is supported by the National Institute of Allergy and Infectious Diseases of the National Institutes of Health grant number K23AI108951.

## Notes

**Conflict of interest:** None of the authors has any financial conflict of interest with the work presented

