## Supplemental Figures for "Inhibition of EZH2 ameliorates lupus-like disease in *MRL/lpr* mice"

**Supplemental Figure 1. Flow gating schematic for human CD19+ B cells, CD14+ monocytes, and CD16b+ neutrophils.** All populations were gated using forward and side scatter and then from singlet cells. Each cell type was then gated based on its representative marker: CD19+ B cells, CD14+ monocytes, and CD16b+ neutrophils.

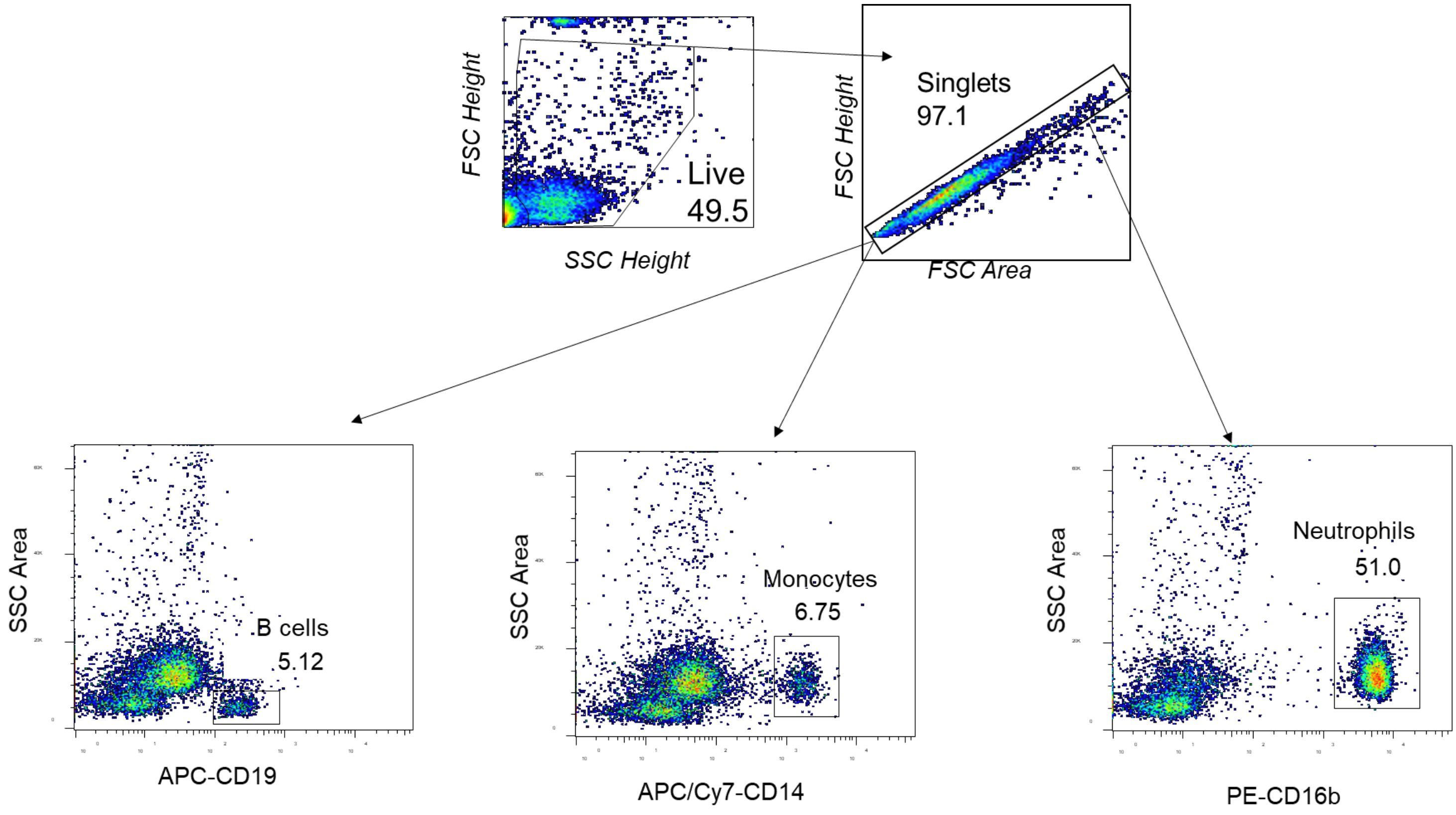

**Supplemental Figure 2. Flow gating schematic for MRL/*lpr* mouse cell types.** All populations were gated using forward and side scatter and then from singlet cells. T cell populations were gated using TCRβ+ cells, and then CD4+, CD8+ or CD4-CD8- for double negative (DN) T cells. B cells were gated using TCRβ- cells and then were identified by CD19+ expression.

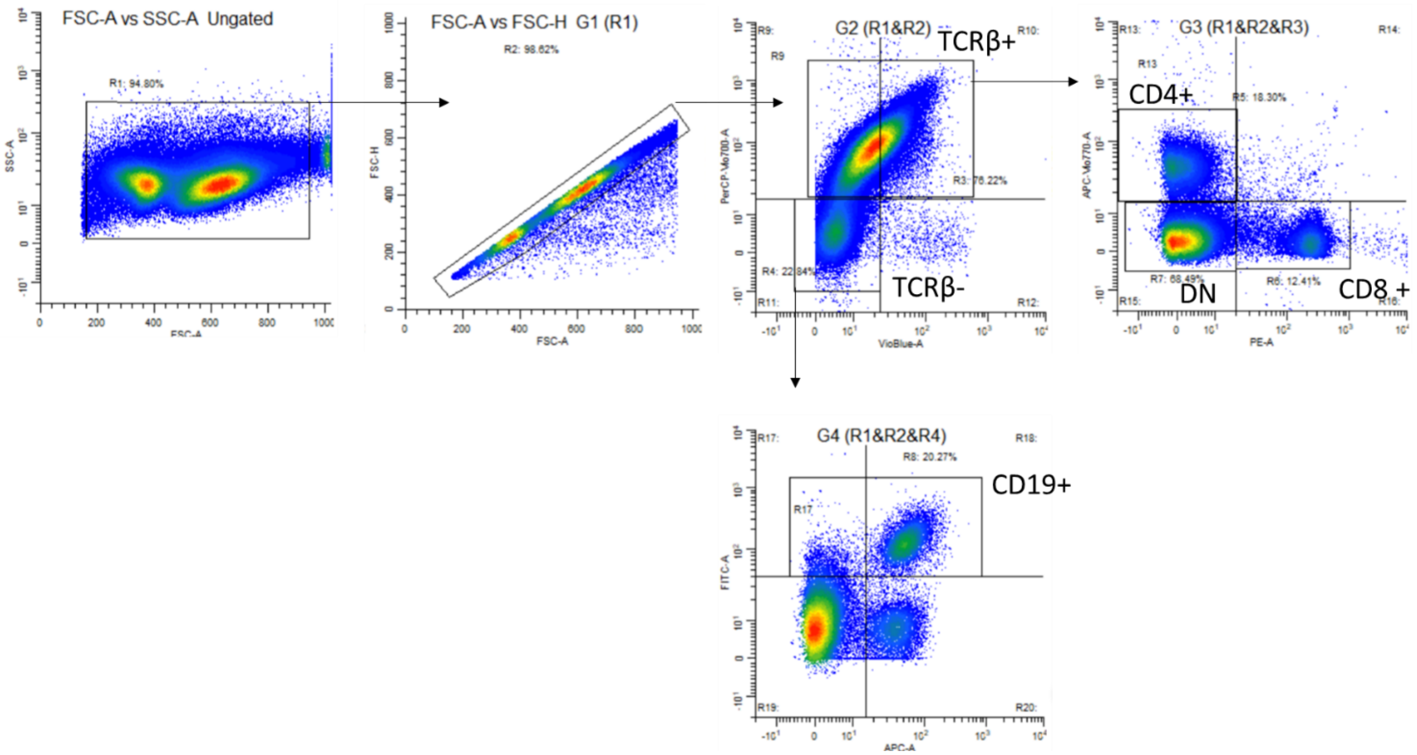

**Supplemental Figure 3. Percentages of glomeruli with no pathologic abnormality.** The percentages of glomeruli with no pathologic abnormality were calculated and differences between DZNep/DZNep and Vehicle/DZNep treatment groups compared to the Vehicle control group were assessed. Comparisons were made using a Kruskal-Wallis with Dunn's multiple comparison test. Results are expressed as mean +/- SEM and $p<0.05$ was considered significant.

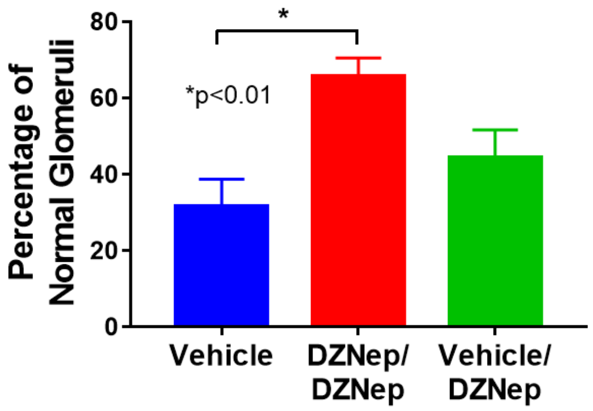

**Supplemental Figure 4. Lymph node weights.** The weights of submaxillary, thoracic, axillary, renal, and mesenteric lymph nodes were significantly decreased for both the DZNep/DZNep and Vehicle/DZNep treatment groups compared to control. Comparisons were made using a Kruskal-Wallis with Dunn's multiple comparison test. Results are expressed as mean +/- SEM and p<0.05 was considered significant.

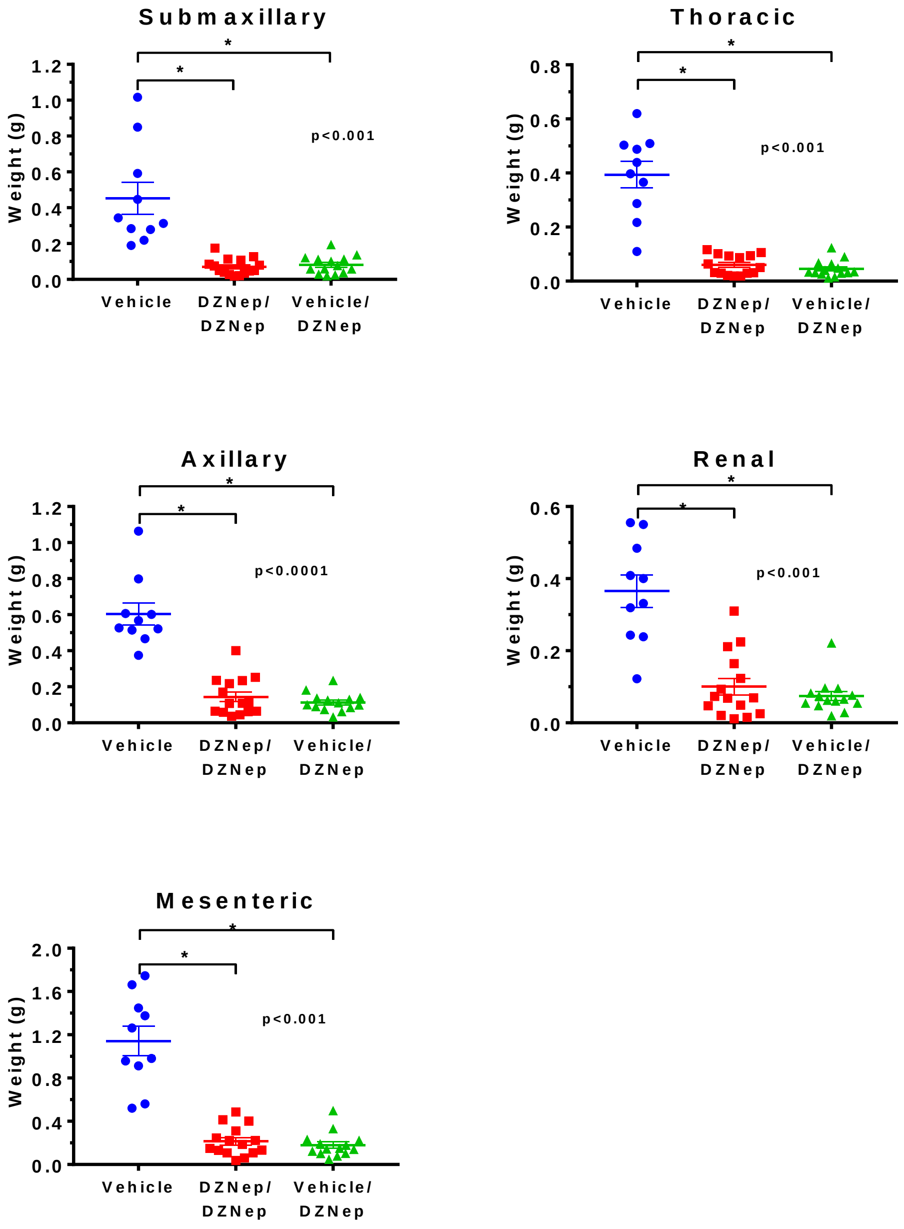
